# A sweet new set of inducible and constitutive promoters for haloarchaea

**DOI:** 10.1101/2023.04.06.535941

**Authors:** Theopi Rados, Katherine Andre, Micaela Cerletti, Alex Bisson

## Abstract

Inducible promoters are one of cellular and molecular biology’s most important technical tools. The ability to deplete, replete, and overexpress genes on demand is the foundation of most functional studies. Here, we developed and characterized a new xylose-responsive promoter (Pxyl), the second inducible promoter system for the model haloarcheon *Haloferax volcanii*. Generating RNA-seq datasets from cultures in the presence of four historically used inducers (arabinose, xylose, maltose, and IPTG), we mapped upregulated genomic regions primarily repressed in the absence of the above inducers. We found a highly upregulated promoter that controls the expression of the *xacEA* (*HVO_B0027-28*) operon in the pHV3 chromosome. To characterize this promoter region, we cloned msfGFP (monomeric superfold green fluorescent protein) under the control of two different 5’ UTR fragments into a modified pTA962 vector: the first 250 bp (P250) and the whole 750 bp intergenic region (P750). The P250 region expressed msfGFP constitutively, and its expression did not respond to the presence or absence of xylose. However, the P750 promoter showed not only to be repressed in the absence of xylose but also expressed higher levels of msfGFP than the previously described inducible promoter PtnaA in the presence of the inducer. Finally, we validated the inducible Pxyl promoter by reproducing morphological phenotypes already described in the literature. By overexpressing the tubulin-like FtsZ1 and FtsZ2, we observed similar but slightly more pronounced morphological defects than the tryptophan-inducible promoter PtnaA. FtsZ1 overexpression created larger, deformed cells, whereas cells overexpressing FtsZ2 were smaller but mostly retained their shape. In summary, this work contributes a new xylose-inducible promoter, Pxyl, that can be used simultaneously with the well-established PtnaA in functional studies in *H. volcanii*.

## Introduction

Inducible promoters have been essential tools for molecular and cell biology studies in bacteria and eukaryotes, allowing for control over the expression of genes of interest in terms of both timing and strength of expression. Among the multiple uses of inducible promoters are dynamic studies of gene expression *in vivo*, timed expression of tagged protein fusions, and depletion assays of essential genes. On the other hand, constitutive promoters have a wide range of applications, from expressing selective marker genes to consistent overexpression of proteins.

Like most model organisms, the budding yeast *Saccharomyces cerevisiae* has a breadth of inducible promoters, most relying on ethanol and sugars (Weinhandl et al., 2014). These have later been replaced by variations of bacterial promoters due to tighter control of the latter. Additionally, there are two kinds of light-induced promoters used in mammalian cell studies, based on a light-oxygen-voltage (LOV) domain from the fungus *Neurospora crassa* and the *Arabidopsis* photoreceptor (Kallunki et al., 2019), which have been used in the research and identification of essential signaling pathways such as mTOR and Hippo (Kallunki et al., 2019). In plants, there is also a variety of available inducible promoters that can be used in *Arabidopsis thaliana* and other model systems, both chemically induced by compounds distinct from those used in mammalian systems (ethanol, dexamethasone, and β-estradiol) (Borghi, 2010) as well as the promoter of the *Arabidopsis* heat-shock protein *HSP18*.*2* gene, which is a strong inducible promoter that is activated by heat shock (Matsuhara et al., 2000; Takahashi and Komeda, 1989). In 2020, a xylose-inducible promoter was first cloned and used to express heterologous proteins in the thermoacidophilic archaeon *Sulfolobus acidocaldarius* (van der Kolk et al., 2020). Currently, the only inducible promoter available in the model haloarchaeaon *Haloferax volcanii* is the tryptophan-inducible PtnaA (Allers et al., 2010).

Proteins isolated from halophiles have been increasingly used in biotechnology, from cosmetics manufacturing to bioremediation (Haque et al., 2020). Methods for expressing and purifying halophilic proteins, which commonly misfold and aggregate when expressed in *Escherichia coli* (Large et al., 2007), have been described in *Haloferax volcanii* (Allers, 2010) using the PtnaA promoter. Overexpression of such proteins for purification could benefit from stronger inducible or strong constitutive promoters being available in *H. volcanii*. Likewise, as *H. volcanii* emerges as a well-studied archaeal model due to its relative ease of cultivation and established genetics (Pohlschroder and Schulze, 2019), the development of molecular biology tools that allow the expression of multiple genes simultaneously becomes essential.

Recently, Nußbaum and coworkers (Nußbaum et al., 2021) reported that experiments on the hierarchy of recruitment of SepF by FtsZ1 or FtsZ2 were not feasible due to a lack of a second inducible promoter in *H. volcanii*. Despite the considerable characterization of the metabolism and catabolism of sugars in *H. volcanii* (Johnsen et al., 2009, 2013, 2015), little is known about how these promoters behave under ectopic expression during live-cell imaging experiments. To address the needs of the *Haloferax* community, we have here characterized two versions of the same promoter region: a strong constitutive and a new xylose-inducible promoter to be used as a tool for genetic studies in *H. volcanii*.

## Methods

### *Haloferax volcanii* cultures

Cells were grown in 16×25 mm glass tubes with 3 ml of Hv-Cab (de Silva et al., 2021) at 42°C under constant agitation with the inducers D-xylose (Thermo Scientific Chemicals), L-arabinose (Thermo Scientific Chemicals), IPTG (Fisher Bioreagents), D-maltose (Fisher Bioreagents), or L-tryptophan (Thermo Scientific Chemicals) in the concentrations indicated.

### Cloning and transformations

The eTR8 vector was constructed by Gibson assembly (Gibson et al., 2009) from two PCR fragments using the oligos oTR26 and oTR27 (msfGFP) and a linearized pTA962 digested with NdeI.

The eAD08 vector was constructed by Gibson assembly from two PCR fragments using the oligos oBL340 and oJM96 (to amplify the msfGFP fragment) and the linearized pTA962 digested with NcoI.

The pAL250::msfGFP and pAL750::msfGFP vectors were constructed by Gibson assembly from three PCR fragments using the oligos oBL169 and oBL345 (msfGFP, eTR8 as a template), oBL343 and oBL344 (the first 250 bp upstream to the *HVO_B0027* start codon using the *H. volcanii* strain DS2 gDNA as a template) or oBL343 and oBL345 (the first 750 bp upstream to the *HVO_B0027* start codon using the *H. olcanii* strain DS2 gDNA as a template), and a linearized pTA962 previously digested with KpnI and BamHI.

The pAL750 vector was created by Gibson assembly from three PCR fragments using the pAL::msfGFP vector as a template and the oligos oZC23 and oTR150 (*pHV2 ori*), oTR151 and oSB33 (pyrE2-Pxyl_750_ region), and oBL97 and oBL105 (*E. coli oriC* and AmpR cassette).

The pAL750::ftsZ1 and pAL750::ftsZ2 vectors were cloned by Gibson assembly of two fragments using the oligos oTR151 and oTR152 (ftsZ1) or oTR153 and oTR154 (ftsZ2) using the *H. volcanii* strain DS2 gDNA as a template, and a linear fragment of the pAL vector digested with NdeI.

All Gibson reactions were transformed into competent *E. coli* DH5α cells and clones confirmed by whole-plasmid sequencing (Plasmidsaurus). Plasmid preps were then transformed into *H. volcanii* using the method previously described (Dyall-Smith M., 2009), with 0.5 M EDTA (Thermo Scientific Catalog #J15694-AE) and PEG600 (Sigma, catalog # 87333-250G-F).

### Growth curves

Cells were grown to an OD_600_ of ∼0.5 and diluted to an OD_600_ of 0.05. Then, 200 μL of culture was placed in a 96-well flat-bottom plate (Corning Inc.). All wells surrounding the plate’s edge (A and H rows, 1 and 12 columns) were filled with 200 μl of water to prevent media evaporation. Growth curves of three biological triplicates were performed using an EPOCH 2 microplate spectrophotometer (Agilent) with constant orbital shaking at 42°C. Data points were collected every 30 minutes.

### Microscopy and Image Analysis

Cells were grown in Hv-Cab or YPC medium (Allers et al., 2010) to mid-exponential or stationary OD_600_ without or with inducers as indicated. Cultures were then concentrated 10-fold by centrifuging (3000xg for 2-5 minutes), and a 3 μL droplet of culture was placed on a 60×24 mm coverslip and gently covered with a 1.5×1.5 cm 0.25% Hv-Cab agarose pad (SeaKem LE Agarose, Lonza). Staining with Ethidium Bromide (Invitrogen, catalog #15585-011) was performed by adding 3 μg/ml of the dye into the cell culture and incubating at 42°C for 5 minutes. Cells were then imaged at 42°C using a Nikon TI-2 Nikon Inverted Microscope within an Okolab H201 enclosure. Phase-contrast and GFP-fluorescence images were acquired with a Hamamatsu ORCA Flash 4.0 v3 sCMOS Camera (6.5 μm/pixel), a CFI PlanApo Lambda 100x DM Ph3 Objective, and a Lumencor Sola II Fluorescent LED (380-760 nm). Image analysis was performed using FIJI (Schindelin et al., 2012). Cells were segmented using the rolling ball background subtraction (value=20) on phase-contrast images, followed by thresholding (default) and using the analyze particles function with a minimum particle size of 0.2 μm^2^. The mask was then used to acquire shape descriptor data and applied to the GFP channel (after background subtraction) for fluorescence quantification.

### RNA extraction and sequencing

All steps were performed at room temperature unless otherwise indicated. Cells were grown to an OD_600_ of ∼0.5 in xylose (10 mM), arabinose (10 mM), IPTG (1 mM), and maltose (1 mM) and harvested by centrifugation (4500xg for 10 minutes). RNA extraction was performed using 1 mL of TriZol (Invitrogen) per sample, followed by vigorous pipetting to lyse all cells in the sample. 200 μL of chloroform was added to the sample mixture, and samples were vortexed for 90 seconds. Samples were then centrifuged at 12,000xg at 4 °C for 15 minutes, and the supernatant was collected and mixed with 2 μL of glycogen (Sigma Aldrich) and 400 μL of isopropanol (Thermo Scientific Chemicals). Samples were incubated overnight at -20 °C. Samples were centrifuged at 21,000xg, 4°C for 30 minutes and washed twice with 75% ethanol (Zaffagni et al., 2022). Samples were treated with DNAse I (NEB) for 12 minutes at 37°C, followed by a second ethanol precipitation. Purified RNA was sent to SeqCenter for ribosome depletion using *Haloferax*specific probes (Table S1) and sequencing. Results were mapped to the *H. volcanii* DS2 genome and analyzed using Geneious 2002.2. Transcripts per million separated by ORF and inducer can be found in Table S2. Raw sequencing data (.fastq and quality control files) is available in Supplemental Materials.

**Table 1:** Plasmids used in this work.

| Alias | Plasmid | Reference |
| --- | --- | --- |
| pTA962 | pTA962 | (Allers et al., 2010) |
| eAD8 | pTA962::PfdX-msfGFP | This work |
| eTR8 | pTA962::msfGFP | This work |
| eTR34 | pAL750::msfGFP | This work |
| eTR36 | pAL750 | This work |
| eTR34 | pAL750::msfGFP | This work |
| eTR38 | pAL750::ftsZ1 | This work |
| eTR40 | pAL750::ftsZ2 | This work |
| eBL82 | pTA962::ftsZ1 | This work |
| eBL86 | pTA962::ftsZ2 | This work |
| eBL227 | pAL250::msfGFP | This work |

**Table 2:**
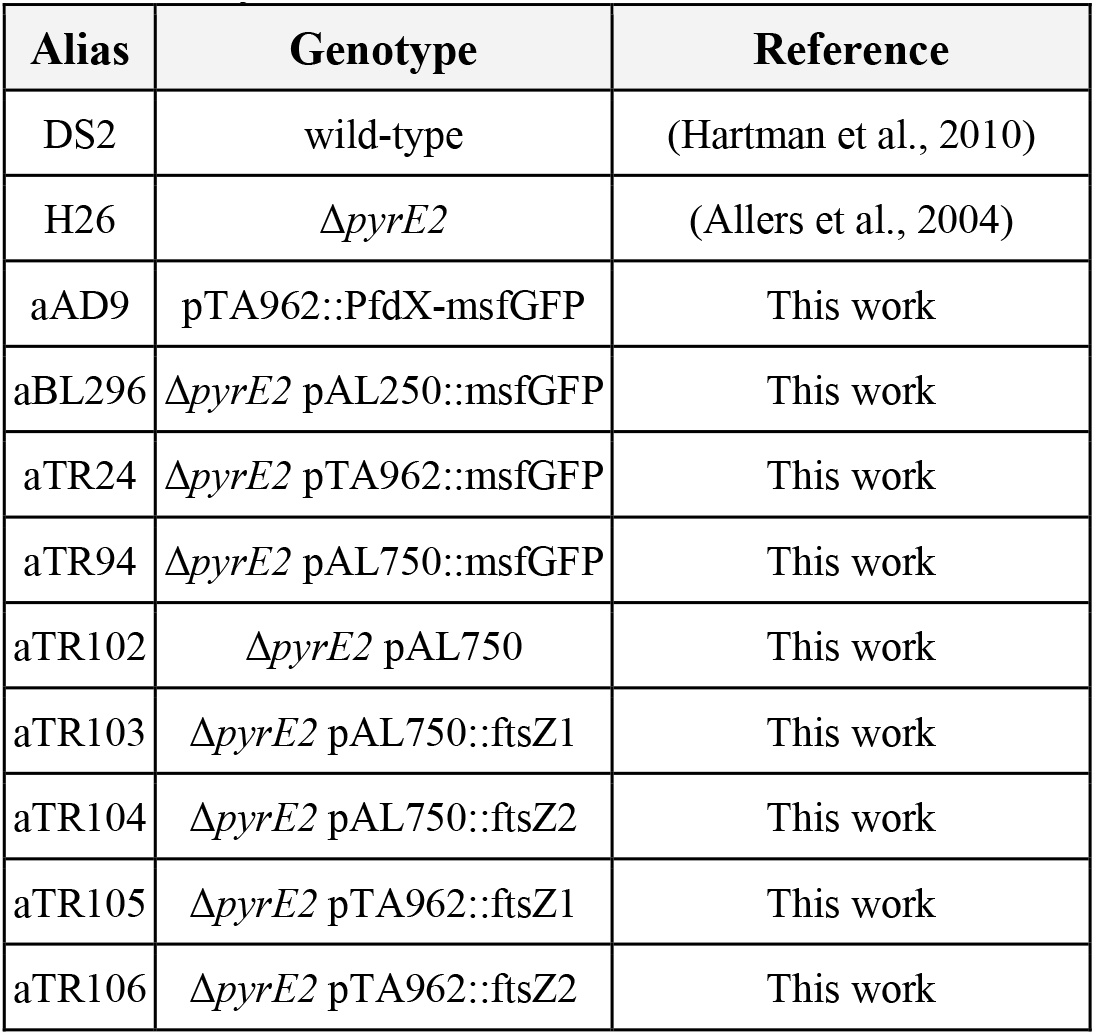
*Haloferax volcanii* strains used in this work.

**Table 3:**
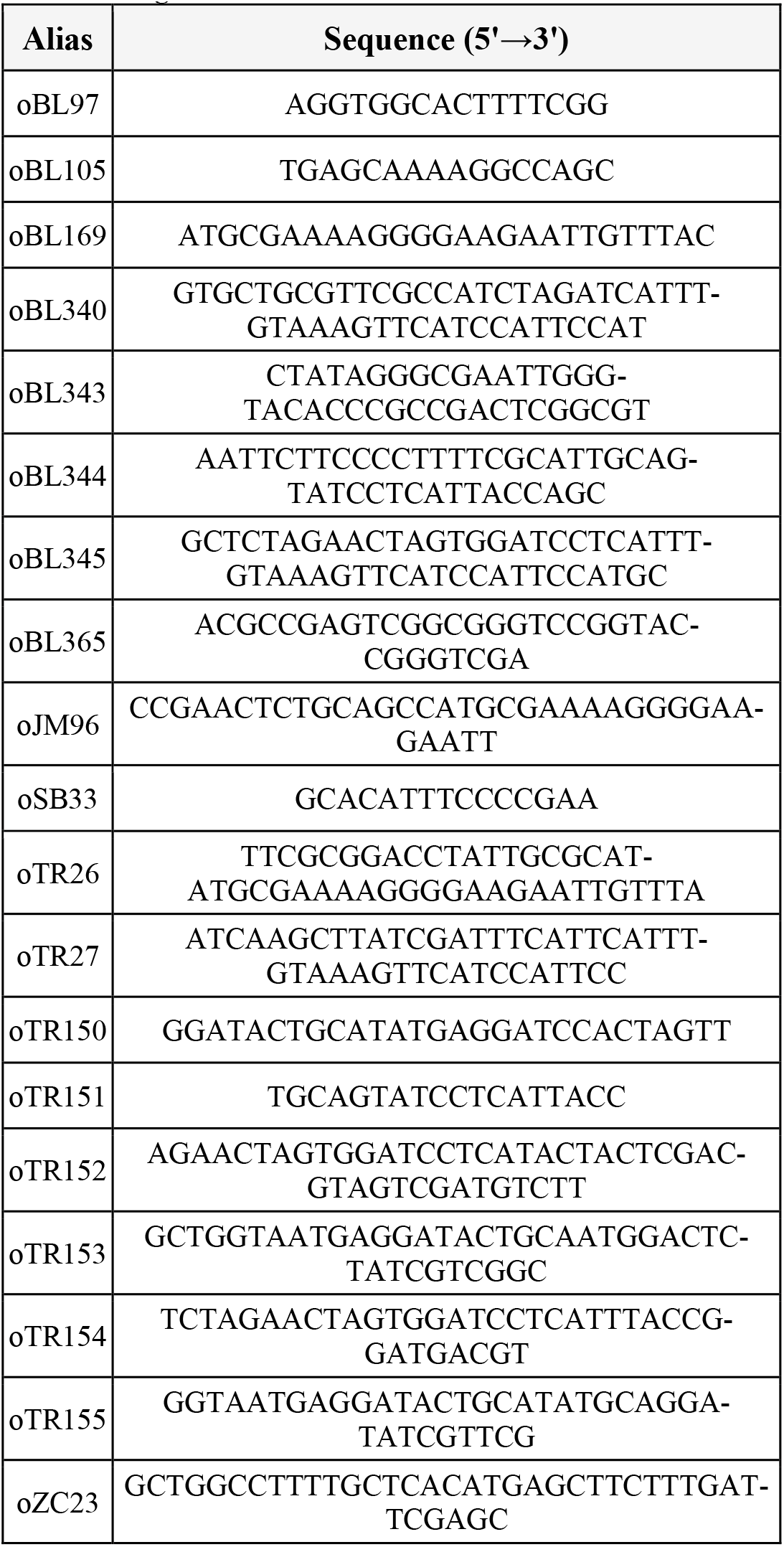
Oligos used in this work.

## Results

### RNA-seq screening to identify new sugar-responsive promoters

To find native inducible promoters, we investigated four different sugars frequently used as inducers in other microbial models: arabinose (Guzman et al., 1995), xylose (Kim et al., 1996), maltose (Ming-Ming et al., 2006), and IPTG (Dubendorff and Studier, 1991). To determine the highest sugar concentrations we could use in *H. volcanii* cultures without compromising growth rates, cell size, and morphology, we titrated each sugar from 1 mM to 100 mM. Based on concentrations used to induce promoters in bacteria, we expected that concentrations higher than 1 mM could yield slow-growing, smaller cells. Surprisingly, we could only observe this outcome from cultures above 10 mM (Fig. 1A-B). Therefore, we focused on concentrations at 10 mM.

**Figure 1:**
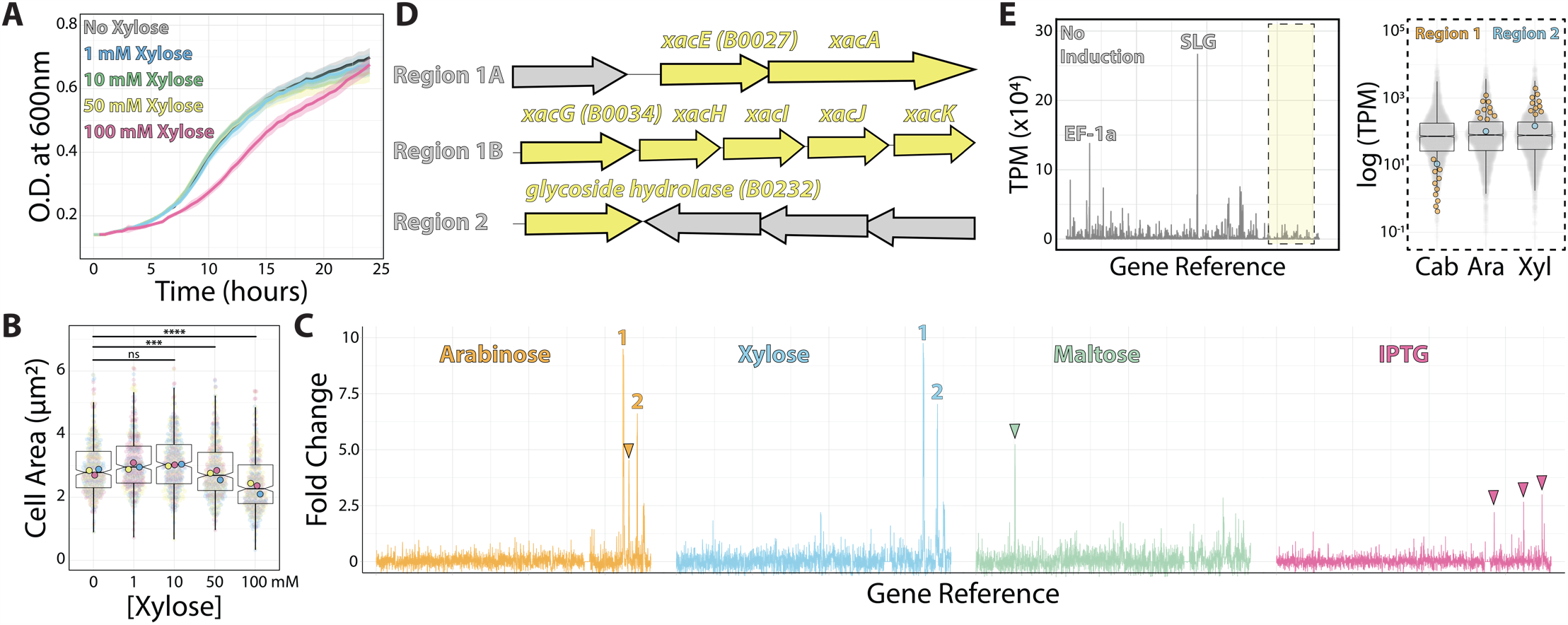
RNA-seq map of inducer-responsive promoters in H. volcanii. (A) growth curves of *H. volcanii* DS2 cells under different xylose concentrations. (B) Cell area measurements at different xylose concentrations by phase contrast microscopy from mid-exponential cultures. (C) Gene expression ratios mapped across the *H. volcanii* genome from RNA-seq datasets of mid-exponential cultures with and without 10 mM arabinose (orange), xylose (blue), maltose (green), and IPTG (pink). Numbers 1 and 2 indicate genomic regions where gene expression increased above 5-fold. Arrowheads indicate promising genomic regions that did not satisfy our arbitrary 5-fold cutoff. (D) locus organization of regions 1 and 2. (E) Expression map (transcripts per million) of genomic regions from RNA-seq datasets without inducers (left) and the relative increase in expression (right) from uninduced (Cab) and arabinose (Ara) and xylose (Xyl).

Next, we employed RNA-seq from mid-exponential cultures with or without each of the four inducers to map candidates for new inducible promoters. Using a set of specific probes for *H. volcanii* (Supplementary Table 1) (Pastor et al., 2022), we obtained a ribosomal RNA depletion of 99.7% from a total number of reads of 120,234,156. By comparing the relative fold change of transcripts per million, we obtained 58 genes from which mRNA levels were significantly upregulated (log_2_ fold change ≥ 1 and p-value ≤ 0.05) for xylose, 42 genes upregulated by maltose, 0 for IPTG, and 45 genes upregulated for arabinose. We identified 15 downregulated genes (log_2_ fold change ≤ -1 and p-value ≤ 0.05) for xylose, 44 genes for maltose, 2 for IPTG, and 10 for arabinose (Supplemental Table 2).

To create a shortlist of promoter candidates we could further validate experimentally, we arbitrarily selected genes above 5-fold or higher ratio change upon the addition of the inducer (Fig. 1C). Above this threshold, we assigned 1 promoter for maltose (*HVO_0562*-*64*), 4 for xylose (*HVO_B0027*-*29, HVO_B0030*-*32, HVO_B0035*, and *HVO_B0036*-*38*), 3 for arabinose (*HVO_B0027*-*29, HVO_B0030*-*32*, and *HVO_B0035)*, and none for IPTG (Fig. 1C and D). In all cases, the expression of candidate genes was upregulated upon the addition of xylose or arabinose (regions 1 and 2). In contrast, the genomic region comprising gene *HVO_B106* did not pass our requirements; it called our attention for being arabinose-specific and not being significantly upregulated under xylose addition (Fig 1C, orange arrowhead). Likewise, genes *HVO_0562-HVO_0566* are specifically upregulated under maltose induction (Fig. 1C, green arrowhead), and three regions (*HVO_A0173, HVO_B0032*, and *HVO_B0303*) respond to IPTG (Fig 1C, pink arrowheads).

Despite observing an approximately 10-fold increase in mRNA levels upon inducer addition, we wanted to ensure these genes were tightly controlled by an inducible promoter with precise linear titration power and not already constitutively expressed at a high basal expression level. To understand the basal expression levels of genes, including regions 1 and 2, we plotted the raw transcription profile (Fig 1E, left panel). We compared it to the transcriptional levels of all other genes (Fig 1E, right panel). As expected, after ribosomal RNA depletion, the transcripts with higher relative values in our datasets were the ones mapping to the S-layer glycoprotein (*csg*, over 26,000 TPM - Transcripts Per Million - across samples) and the translational factor *EF1a* transcripts (above 10,000 TPM across samples). On the other hand, both promoter regions 1 and 2 transcriptional levels are placed in the low quartile range of our dataset in the uninduced sample (1.44 TPM and 23.65 TPM, respectively).

Altogether, genomic regions 1 and 2 are promising candidates for new xylose- and arabinose-inducible promoters for *H. volcanii*. For the context of this work, and based on the dynamic range suggested by our RNA-seq dataset above, we focused on the characterization of the promoter regulating the gene *xacE* (*HVO_B0027*), or genomic region 1, under the induction of xylose.

### The *xacEA* 5’ untranslated region is required for *xacE* repression in the absence of xylose

To test the *xacE* promoter region, we sub-cloned the fluorescent protein msfGFP under the control of two different putative promoter regions: 250 bp (P250) and 750 bp (P750) upstream to *xacE* (Fig. 2A). Plasmids were then transformed into *H. volcanii* H26 (Δ*pyrE2*), and selected on plates without uracil. As a negative control, we used the H26 strain transformed with the empty pAL750 vector (labeled from now on as wild type). As a comparison, we analyzed the expression of msfGFP under the control of the popular inducible promoter PtnaA, which activates upon tryptophan addition (Allers et al., 2010). As a benchmark, we also inserted msfGFP under the control of PfdX, a strong and constitutive promoter used to express the *pyrE2* gene in our vectors as a selective transformation marker (PfdX-msfGFP-pyrE2 operon).

**Figure 2:**
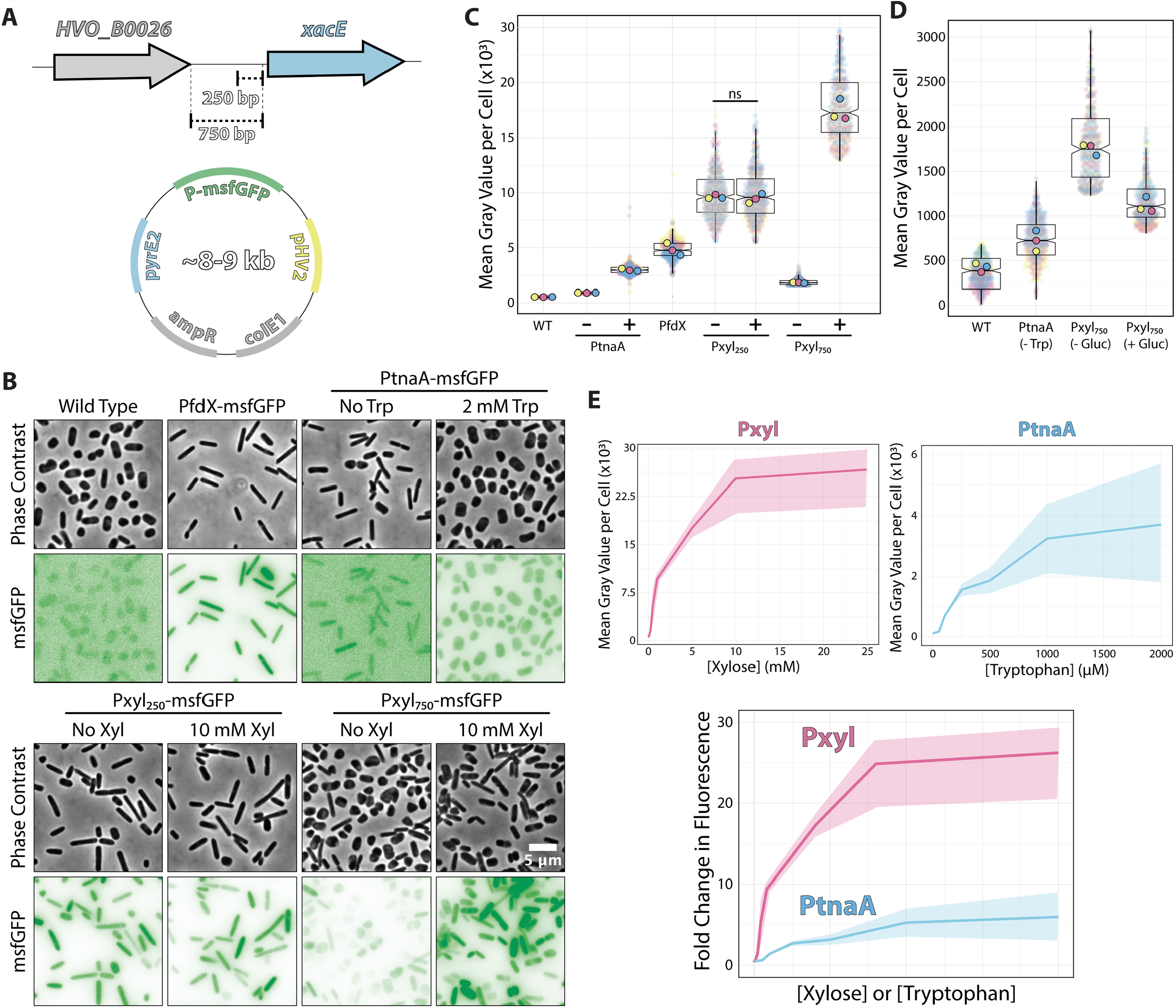
Pxyl constructs can be either strong inducible or constitutive promoters. (A) Fragments of the 5’ intergenic region of *xacE* were used to clone into pAL vectors and map the P250 and P750 promoters tested in this work. (B) Phase-contrast and epifluorescence images of different constructs expressing the msfGFP fluorescent protein. (C) Mean msfGFP fluorescence measurements per cell from images shown in panel B. Each replicate mean and data points are independently labeled with different colors (pink, yellow and blue) (D) Comparative background expression from constructs with and without glucose repression. (E) The dynamic range of PtnaA and Pxyl promoters across different inducer concentrations in raw numbers (top) and normalized (bottom) fold-changes. Shades indicate the 95% confidence interval from triplicate datasets.

Imaging of live cells by phase-contrast and epifluorescence microscopy showed cytoplasmic signal emitted by the msfGFP fluorescent protein from single cells (Fig. 2B). Cells carrying empty plasmids (Fig. 2B, first column) showed low auto-fluorescence at 488 nm excitation compared to cells carrying vectors inducing msfGFP under the control of PfdX and Pxyl promoters. For a quantitative picture of the induction power of each promoter, we segmented individual cells and measured the mean fluorescence per cell from three biological triplicates and graphed using SuperPlots (Goedhart, 2021; Lord et al., 2020). Surprisingly, the P250 promoter was not only insensitive to xylose but also constitutively expressed 2-fold above the PfdX control (10,048±2,256 and 4,967±929 AU, respectively) (Fig. 2C).

In contrast, the P750 promoter harboring the whole 5’ UTR sequence showed approximately a 5.5-fold repression (1,827±848) without induction in comparison to P250. However, msfGFP levels of non-induced P750 cells were still relatively high, 5.7-fold higher than uninduced PtnaA cells (320±14) (Fig. 2B and C). To minimize the transcriptional leakage from P750, we tested whether *Haloferax* cells would present catabolite repression upon adding glucose. This strategy has been successful in various bacterial and yeast systems (Deutscher, 2008; Gancedo, 1998). Adding 20 mM glucose to cultures decreased leakage of the Pxyl promoter, but msfGFP intensity is still 2.7-fold higher than PtnaA cells (Fig. 2D).

However, different from the P250 promoter, the addition of 10 mM xylose to P750 cells resulted in a 10.3-fold increase (18792±4129) in fluorescence intensity with a wider heterogeneity across the population compared to P250. Nevertheless, the unusual decrease in msfGFP expression observed between P250 and P750 suggests the 5’ region of *xacE* may be regulated either at the promoter or mRNA levels.

We also inspected the dynamic range of our P750 promoter compared to PtnaA. By titrating xylose (0 to 25 μM) and tryptophan (0 to 2 mM), we observed a significant improvement from an 11.2-fold linear range for the PtnaA promoter to a 23-fold for the P750 promoter (Fig. 2E). Providing the relatively higher leakage of the P750 promoter levels compared to PtnaA, we concluded that this new construct is a valuable tool for titration experiments and overexpression at high protein levels for functional studies and protein purification directly from *H. volcanii*. Notably, P250 can be used simultaneously with the above inducible promoters as another constitutive promoter in the *Haloferax* community.

Furthermore, we tested the ability to induce the P750 promoter in cells growing in Hv-YPC, a rich medium based on yeast extract instead of casamino acids (Allers et al., 2010). In contrast with Hv-Cab, cells at mid-exponential growth phase showed lower levels of promoter leakage (222±188), 3.6-fold higher than in Hv-Cab (Fig. 3). However, together with the tighter expression of P750, induced cells in Hv-YPC expressed msfGFP at 7.6 times lower than in Hv-Cab (1636±1258), possibly due to catabolic repression in response to a component in yeast extract.

**Figure 3:**
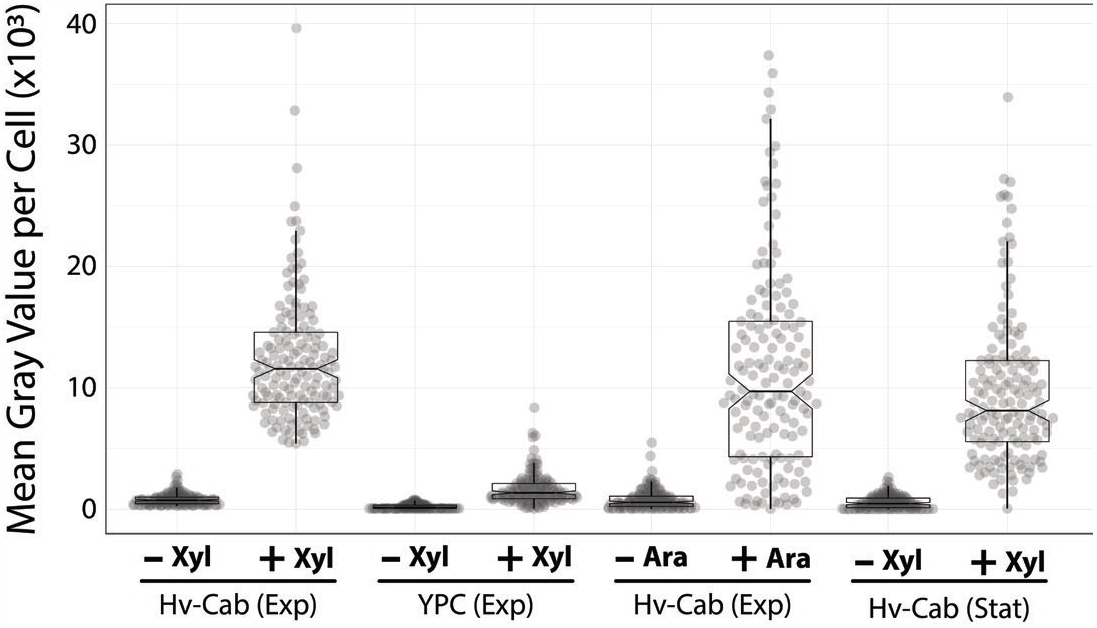
P750 shows different expression profiles in different media, at stationary phase and upon addition of arabinose. Epifluorescence microscopy quantification from exponential growing cells in Hv-Cab medium and Hv-YPC medium induced with 10 mM xylose (first and second plots). Expression of msfGFP was also recorded in cells grown to exponential phase in Hv-Cab medium with 10 mM arabinose (third plot). Cells grown in Hv-Cab to stationary phase and induced with 10 mM xylose (fourth plot).

In addition to Hv-YPC, we also measured the expression profile of P750 at stationary growth phase, and upon the addition of arabinose instead of xylose (Fig. 3). While we observed a similar induction profile from cells in stationary phase and under arabinose induction, both cases showed a wider signal distribution (9851±617 and 11699±797, respectively) compared to xylose-based induction during exponential growth (12395±800).

### Overexpression of the tubulins FtsZ1 and FtsZ2 confirms reported morphological phenotypes

Recently, Liao and colleagues reported the role of two tubulin paralogs (FtsZ1 and FtsZ2) in *H. volcanii*’s cell division (Liao et al., 2021). The authors used PtnaA-controlled overexpression of FtsZ1 and FtsZ2 independently and observed distinct, specific morphological phenotypes related to the overexpression of each paralog. Cells under PtnaA-FtsZ1 overexpression were slightly larger but significantly misshaped compared to the control, whereas PtnaA-FtsZ2 cells looked significantly smaller but showed a more consistent morphology.

To confirm if those phenotypes are reproducible or even enhanced in our new Pxyl system, we cloned *ftsZ1* and *ftsZ2* in the pAL vector and analyzed the cell size and circularity compared to PtnaA-induced cells. Cells overexpressing *ftsZ2* under the Pxyl promoter are slightly smaller (1.2-fold decrease in average cell area) than cells overexpressing *ftsZ2* under the tryptophan-inducible PtnaA (Fig 4A). Meanwhile, cells overexpressing *ftsZ1* under the Pxyl promoter seemed to have more drastic phenotypes than those overexpressing *ftsZ1* using PtnaA, with deformed and enlarged cells (Fig 4A and C). Curiously, a fraction of the population exhibits narrow “cell bridges’’ connecting two enlarged cells (Fig. 4B), similar to those previously described (Rosenshine et al., 1989; Sivabalasarma et al., 2021; von Kügelgen et al., 2021). This indicates that, in our samples, the cell-to-cell bridges observed in phase contrast could be formed by a product of incomplete cell division.

**Figure 4:**
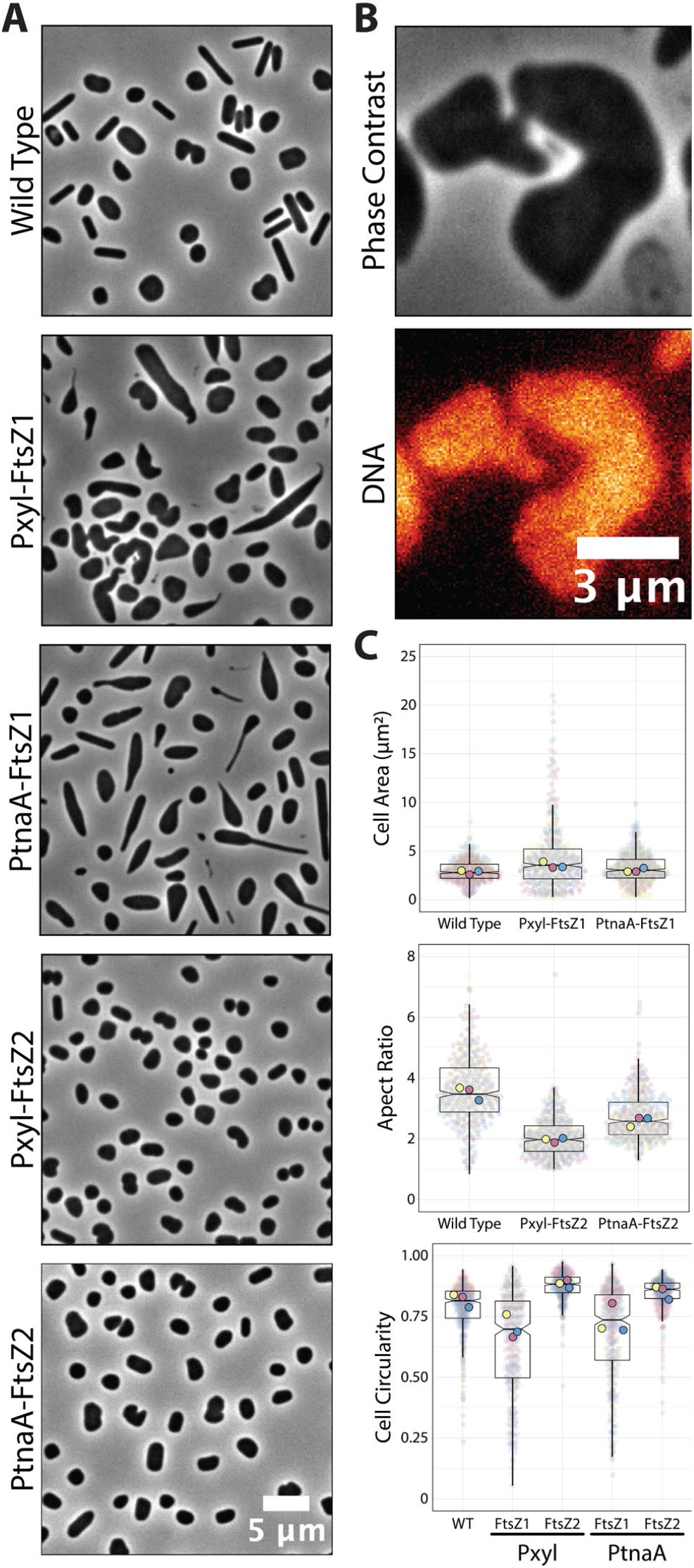
Overexpression of ftsZ1 and ftsZ2 under the Pxyl promoter. (A) Phase-contrast microscopy showing wild-type, overexpressing ftsZ1 or ftsZ2 cells under the Pxyl (5 mM xylose) or PtnaA (2 mM tryptophan) promoters. (B) Representative cell bridge phase-contrast and epifluorescence images. DNA was stained with ethidium bromide. (C) Cell area and circularity measurements from cells overexpressing ftsZ1 and ftsZ2 with the Pxyl (5 mM xylose) and the PtnaA (2 mM tryptophan) promoters. Each replicate mean and data points are independently labeled with different colors (pink, yellow and blue).

## Discussion

Inducible promoters have been an invaluable resource in basic molecular biology research for the past 60 years, since the early days of the “PaJaMa Experiments” (Lewis, 2011). The addition of xylose to the culture medium was first shown to induce the expression of genes in *E. coli* (Batt et al., 1985) and *B. subtilis* in 1988 (Gärtner et al., 1988). This report describes the characterization of a new xylose-inducible promoter for the halophilic archaeon *H. volcanii*, a well-studied archaeal model (Pohlschroder and Schulze, 2019).

Using RNA-seq in cultures growing with and without four different inducers, we identified multiple promoter regions in which transcript levels were upregulated upon the addition of arabinose (total of 23 promoters, 3 above the arbitrary cut-off), xylose (total of 18 promoters, 4 above the arbitrary cut-off), maltose (total of 6 promoters, none above the arbitrary cut-off) and IPTG (total of 2 promoters, none above the arbitrary cut-off). Interestingly, our most promising promoter chosen to be characterized was already noted in past studies using DNA microarrays and C13 isotope tracking (Johnsen et al., 2009) and further characterized *in vitro* and *in vivo* (Johnsen et al., 2015, 2013). However, their measurements (ranging from 100- to 400-fold increase upon arabinose addition) were greatly overestimated compared to our observations from RNA-seq (9.8-fold, Fig. 1C) and live-cell microscopy (10.3-fold increase, Fig. 2C). Curiously, against the anecdotal knowledge among researchers in the field, the PtnaA promoter showed a relatively low leakage in cultures without tryptophan (Fig. 2D).

One interesting previously unmentioned feature of the transcriptional regulation of the *xacEA* operon is that the P750 promoter not only is significantly repressed in the absence of xylose, but P750 shows a 1.9-fold increase in msfGFP signal in comparison to the constitutive P250 promoter (Fig. 2C). The mechanistic details are still elusive, but it is possible that the extreme 5’ UTR is the target of transcriptional factors competing to repress and activate the expression of *xacEA*. A good candidate for an activator is XacR (HVO_B0040), an IclR transcriptional factor family shown to moonlight between repression and activation (Krell et al., 2006; Pan et al., 2011). In agreement with previous observations in bacteria, XacR in *H. volcanii* was shown to be required to activate *xacE* expression *in vivo* (Johnsen et al., 2015).

We have shown that, through higher levels of expression than with the available promoter PtnaA, cells overexpressing the tubulin-like FtsZ1 had subtle but clear morphology defects (Fig. 4) beyond those previously described (Liao et al., 2021). Interestingly, FtsZ2 overexpression under our Pxyl system did not result in a convincing change to PtnaA-FtsZ2 cells, in agreement with data suggesting that FtsZ2 proteins are more unstable compared to FtsZ1 (Liao et al., 2021). Alternatively, the FtsZ2 function could be coupled with other factors that are more limited in the cell, and therefore a higher concentration of FtsZ2 would not linearly scale with cell size.

Aside from the Pxyl promoter, we mapped other promoter regions that showed to be independently upregulated upon the addition of maltose and IPTG (Fig. 1C). Future characterization of these promoters may expand the toolbox of inducible promoters in *H. volcanii*.

## Acknowledgments

We thank the Bisson Lab, Kiwi Shaw-Dodge, and Rosana De Castro (Universidad Nacional de Mar del Plata, Mar del Plata) for their insightful comments on our manuscript. We appreciate Sebastian Kadener and Sinead Nguyen (Brandeis University) for their help in optimizing the total RNA extraction from *H. volcanii*. We also thank Amy Schmid and Mar Martinez-Pastor (Duke University) for sharing data before publication on their ribodepletion probes. This work was supported by the Human Frontiers Science Program funding (RGY0074/2021) and Life Sciences-Moore–Simons Project on the Origin of the Eukaryotic Cell (https://doi.org/10.46714/735929LPI) awarded to AB. AB is a Pew Scholar in the Biomedical Sciences, supported by The Pew Charitable Trusts. MC was supported by the CONICET Partial Financing Program for Stays Abroad for Assistant Researchers.

## Author Contributions

AB and TR conceived the study. AB, MC, TR, and KA designed the experiments. TR and KA performed the experiments and analyzed the data. AB and TR wrote the first draft. All authors reviewed and approved the submitted version.

## Data Availability

The complete raw RNA-seq datasets presented in this study can be found in online repositories: PRJNA953041 (no induction), PRJNA953037 (xylose), PRJNA953035 (IPTG), PRJNA953033 (Arabinose), PRJNA953034 (Maltose).

## References

Allers, T., 2010. Overexpression and purification of halophilic proteins in Haloferax volcanii. Bioeng. Bugs 1, 290–292. https://doi.org/10.4161/bbug.1.4.11794

Allers, T., Barak, S., Liddell, S., Wardell, K., Mevarech, M., 2010. Improved strains and plasmid vectors for conditional overexpression of His-tagged proteins in Haloferax volcanii. Appl. Environ. Microbiol. 76, 1759–69. https://doi.org/10.1128/AEM.02670-09

Allers, T., Ngo, H.-P., Mevarech, M., Lloyd, R.G., 2004. Development of additional selectable markers for the halophilic archaeon Haloferax volcanii based on the leuB and trpA genes. Appl. Environ. Microbiol. 70, 943–53. https://doi.org/10.1128/aem.70.2.943-953.2004

Batt, C.A., Bodis, M.S., Picataggio, S.K., Claps, M.C., Jamas, S., Sinskey, A.J., 1985. Analysis of xylose operon regulation by Mud (Apr, lac) fusion: trans effect of plasmid coded xylose operon. Can. J. Microbiol. 31, 930–933. https://doi.org/10.1139/m85-174

Borghi, L., 2010. Inducible Gene Expression Systems for Plants, in: Hennig, L., Köhler, C. (Eds.), Plant Developmental Biology: Methods and Protocols, Methods in Molecular Biology. Humana Press, Totowa, NJ, pp. 65–75. https://doi.org/10.1007/978-1-60761-765-5_5

de Silva, R.T., Abdul-Halim, M.F., Pittrich, D.A., Brown, H.J., Pohlschroder, M., Duggin, I.G., 2021. Improved growth and morphological plasticity of Haloferax volcanii. Microbiology 167, 001012. https://doi.org/10.1099/mic.0.001012

Deutscher, J., 2008. The mechanisms of carbon catabolite repression in bacteria. Curr. Opin. Microbiol. 11, 87–93. https://doi.org/10.1016/j.mib.2008.02.007

Dubendorff, J.W., Studier, F.W., 1991. Controlling basal expression in an inducible T7 expression system by blocking the target T7 promoter with lac repressor. J. Mol. Biol. 219, 45– 59. https://doi.org/10.1016/0022-2836(91)90856-2

Dyall-Smith M., 2009. The Halohandbook: Protocols for Halobacterial Genetics Ver 7.2.

Gancedo, J.M., 1998. Yeast Carbon Catabolite Repression. Microbiol. Mol. Biol. Rev. 62, 334–361.

Gärtner, D., Geissendörfer, M., Hillen, W., 1988. Expression of the Bacillus subtilis xyl operon is repressed at the level of transcription and is induced by xylose. J. Bacteriol. 170, 3102–3109.

Gibson, D.G., Young, L., Chuang, R.-Y., Venter, J.C., Hutchison, C.A., Smith, H.O., 2009. Enzymatic assembly of DNA molecules up to several hundred kilobases. Nat. Methods 6, 343– 5. https://doi.org/10.1038/nmeth.1318

Goedhart, J., 2021. SuperPlotsOfData—a web app for the transparent display and quantitative comparison of continuous data from different conditions. Mol. Biol. Cell 32, 470–474. https://doi.org/10.1091/mbc.E20-09-0583

Guzman, L.M., Belin, D., Carson, M.J., Beckwith, J., 1995. Tight regulation, modulation, and high-level expression by vectors containing the arabinose PBAD promoter. J. Bacteriol. 177, 4121–4130.

Haque, R.U., Paradisi, F., Allers, T., 2020. Haloferax volcanii for biotechnology applications: challenges, current state and perspectives. Appl. Microbiol. Biotechnol. 104, 1371–1382. https://doi.org/10.1007/s00253-019-10314-2

Hartman, A.L., Norais, C., Badger, J.H., Delmas, S., Haldenby, S., Madupu, R., Robinson, J., Khouri, H., Ren, Q., Lowe, T.M., Maupin-Furlow, J., Pohlschroder, M., Daniels, C., Pfeiffer, F., Allers, T., Eisen, J.A., 2010. The complete genome sequence of Haloferax volcanii DS2, a model archaeon. PloS One 5, e9605. https://doi.org/10.1371/jour-nal.pone.0009605

Johnsen, U., Dambeck, M., Zaiss, H., Fuhrer, T., Soppa, J., Sauer, U., Schönheit, P., 2009. d-Xylose Degradation Pathway in the Halophilic Archaeon Haloferax volcanii. J. Biol. Chem. 284, 27290–27303. https://doi.org/10.1074/jbc.M109.003814

Johnsen, U., Sutter, J.-M., Schulz, A.-C., Tästensen, J.-B., Schönheit, P., 2015. XacR – a novel transcriptional regulator of D-xylose and L-arabinose catabolism in the haloarchaeon Haloferax volcanii. Environ. Microbiol. 17, 1663–1676. https://doi.org/10.1111/1462-2920.12603

Johnsen, U., Sutter, J.-M., Zaiß, H., Schönheit, P., 2013. l-Arabinose degradation pathway in the haloarchaeon Haloferax volcanii involves a novel type of l-arabinose dehydrogenase. Extremophiles 17, 897–909. https://doi.org/10.1007/s00792-013-0572-2

Kallunki, T., Barisic, M., Jäättelä, M., Liu, B., 2019. How to Choose the Right Inducible Gene Expression System for Mammalian Studies? Cells 8, 796. https://doi.org/10.3390/cells8080796

Kim, L., Mogk, A., Schumann, W., 1996. A xylose-inducible Bacillus subtilis integration vector and its application. Gene 181, 71–76. https://doi.org/10.1016/s0378-1119(96)00466-0

Krell, T., Molina-Henares, A.J., Ramos, J.L., 2006. The IclR family of transcriptional activators and repressors can be defined by a single profile. Protein Sci. Publ. Protein Soc. 15, 1207–1213. https://doi.org/10.1110/ps.051857206

Large, A., Stamme, C., Lange, C., Duan, Z., Allers, T., Soppa, J., Lund, P.A., 2007. Characterization of a tightly controlled promoter of the halophilic archaeon Haloferax volcanii and its use in the analysis of the essential cct1 gene. Mol. Microbiol. 66, 1092–1106. https://doi.org/10.1111/j.1365-2958.2007.05980.x

Lewis, M., 2011. A tale of two repressors – a historical perspective. J. Mol. Biol. 409, 14–27. https://doi.org/10.1016/j.jmb.2011.02.023

Liao, Y., Ithurbide, S., Evenhuis, C., Löwe, J., Duggin, I.G., 2021. Cell division in the archaeon Haloferax volcanii relies on two FtsZ proteins with distinct functions in division ring assembly and constriction. Nat. Microbiol. 6, 594–605. https://doi.org/10.1038/s41564-021-00894-z

Lord, S.J., Velle, K.B., Mullins, R.D., Fritz-Laylin, L.K., 2020. SuperPlots: Communicating reproducibility and variability in cell biology. J. Cell Biol. 219, e202001064. https://doi.org/10.1083/jcb.202001064

Matsuhara, S., Jingu, F., Takahashi, T., Komeda, Y., 2000. Heatshock tagging: a simple method for expression and isolation of plant genome DNA flanked by T-DNA insertions. Plant J. 22, 79–86. https://doi.org/10.1046/j.1365-313x.2000.00716.x

Ming-Ming, Y., Wei-Wei, Z., Xi-Feng, Z., Pei-Lin, C., 2006. Construction and characterization of a novel maltose inducible expression vector in Bacillus subtilis. Biotechnol. Lett. 28, 1713–1718. https://doi.org/10.1007/s10529-006-9146-z

Nußbaum, P., Gerstner, M., Dingethal, M., Erb, C., Albers, S.-V., 2021. The archaeal protein SepF is essential for cell division in Haloferax volcanii. Nat. Commun. 12, 3469. https://doi.org/10.1038/s41467-021-23686-9

Pan, Y., Fiscus, V., Meng, W., Zheng, Z., Zhang, L.-H., Fuqua, C., Chen, L., 2011. The Agrobacterium tumefaciens Transcription Factor BlcR Is Regulated via Oligomerization. J. Biol. Chem. 286, 20431–20440. https://doi.org/10.1074/jbc.M110.196154

Pastor, M.M., Sakrikar, S., Rodriguez, D.N., Schmid, A.K., 2022. Comparative Analysis of rRNA Removal Methods for RNA-Seq Differential Expression in Halophilic Archaea. Biomolecules 12, 682. https://doi.org/10.3390/biom12050682

Pohlschroder, M., Schulze, S., 2019. Haloferax volcanii. Trends Microbiol. 27, 86–87. https://doi.org/10.1016/j.tim.2018.10.004

Rosenshine, I., Tchelet, R., Mevarech, M., 1989. The mechanism of DNA transfer in the mating system of an archaebacterium. Science 245, 1387–9.

Schindelin, J., Arganda-Carreras, I., Frise, E., Kaynig, V., Longair, M., Pietzsch, T., Preibisch, S., Rueden, C., Saalfeld, S., Schmid, B., Tinevez, J.-Y.J.-Y., White, D.J., Hartenstein, V., Eliceiri, K., Tomancak, P., Cardona, A., Liceiri, K., Tomancak, P. A. C., 2012. Fiji: an open source platform for biological image analysis. Nat. Methods 9, 676–682. https://doi.org/10.1038/nmeth.2019.Fiji

Sivabalasarma, S., Wetzel, H., Nußbaum, P., van der Does, C., Beeby, M., Albers, S.-V., 2021. Analysis of Cell–Cell Bridges in Haloferax volcanii Using Electron Cryo-Tomography Reveal a Continuous Cytoplasm and S-Layer. Front. Microbiol. 11.

Takahashi, T., Komeda, Y., 1989. Characterization of two genes encoding small heat-shock proteins in Arabidopsis thaliana. Mol. Gen. Genet. MGG 219, 365–372. https://doi.org/10.1007/BF00259608

van der Kolk, N., Wagner, A., Wagner, M., Waßmer, B., Siebers, B., Albers, S.-V., 2020. Identification of XylR, the Activator of Arabinose/Xylose Inducible Regulon in Sulfolobus acidocaldarius and Its Application for Homologous Protein Expression. Front. Microbiol. 11, 1066. https://doi.org/10.3389/fmicb.2020.01066

von Kügelgen, A., Alva, V., Bharat, T.A.M., 2021. Complete atomic structure of a native archaeal cell surface. Cell Rep. 37, 110052. https://doi.org/10.1016/j.celrep.2021.110052

Weinhandl, K., Winkler, M., Glieder, A., Camattari, A., 2014. Carbon source dependent promoters in yeasts. Microb. Cell Factories 13, 5. https://doi.org/10.1186/1475-2859-13-5

Zaffagni, M., Harris, J.M., Patop, I.L., Pamudurti, N.R., Nguyen, S., Kadener, S., 2022. SARS-CoV-2 Nsp14 mediates the effects of viral infection on the host cell transcriptome. eLife 11, e71945. https://doi.org/10.7554/eLife.71945

