## Supplementary material for "A sweet new set of inducible and constitutive promoters for haloarchaea": Table S1

Table S1. *Haloferax volcanii*-specific ribodepletion probes.

Sequence

CCTACCCCGGACGTACCCCTCGAGCCTTCCGGTCCCAAGCCACGTGGTTT

ATCGTCATCACTGTAGTCGGAGCTGGTGAGATGTCCGGCGTTGAGTCCAA

GTTCGTTTTTCAAAACGTACGACGTAACATCGGCTTCACTCGAGTCATAC

GACCTACCGTTGCCCGTTCCTTCCTCCGCTTTAGCAGCGGCAGTCCTTCT

GCACCGCCCTGTTCGGGGTGCTTTTCAGCGTTCGCTCACGCTACTTGTTC

TACGACGTTCGTAAGTTCGGAGTTTGACAGGGAAGCGAACTCCTCTCGGA

CCTACTAAGATGTTTCAATTCGGAACGTTCCCCATTGCGCGAGGCAATTG

CTACGGAATCGTGTCATAAGACACGCCGCCAGGTCTCGGAGGTGGATTTG

ACCACGTCTCAGCGACAGCCTGCAGTAAAGCTCTATAGGGTCTTCGCTTC

AGGAGGTGATCCAGCCGCAGATTCCCCTACGGCTACCTTGTTACGACTTA

TCGTTTACAGCTAGGACTACCCGGGTATCTAATCCGGTTCGAGACCCTAG

GTTTCGGGTCGCATCCGGTTGGCTCCCCGCCCTTGAAGACGGTGGCCCTG

CCCGCTCGGTGTTTCCTAAAGCGTCGACGTTTGGTGCAAGAAGGTACTGG

GTAGCAAGCCACCGGGTCGATATGTGCTCTTGCCGGTGACGACTCTGTTA

CTGCCTCGGTGGCCCGCTGGCGCGGGTACCTGTGTAATGGCTCCTACCTA

TTCTGTCATCTACGGGTCCCATGAATGAACCTCGTAGGTTCGCTAGACCA

GTGCAAGGAGCAGGGACGTATTCACCGCACTCTATTGAAATGCGATTACT

AGTCTATCGAACTCGTCTTCTACGAGTGTCCTCAGTGGAACTTCTTTTCC

CACCGTTCGTTCGCTTCGTGCCGTTACGGCTTCCACGAATTTCGGTGATT

GCCCGGCTACCGGTGTCCTCCGCCAGGAGTGAGAGTCGCAGTCACTAACG

GATCTCGGTACGAGCTTCTTGCTTGTCTTTTCACGGGCTCTAGGTACCGC

GTCGGATCCGTCTTCTCGAGGTGCTTTCGCCATCGGTGGTCCGTCCAGGA

GCTGTTGCCAAGTTCGATGGGCCTTTCACCCCTACACCAGAGTCACGAGA

TCGAAACTGCTGCTGGCAATTAGAAGTGCGGGTCTCGCTCGTTGCCTGAC

AATAGGCGAACAACCTCACCCTTGCCTGCTTCTGCACAGGCAGGATGGAG

CGTATCGGTTTCCCTATGCCTCCGGGGGTTCACCCCTTAGACTCGCCAGT

AGACTCCGCACTGGAACGTACTGTTCACCGGGCCCAACGTTGGGACAGTG

GAATGTGGCTTGGACTGTTAGTGCTCGCGGGCTTAACAACTCGTTACCTC

CAGTTGAGCTGGCGGATTTCCCGATGAACCTTCGTGGCCGGCTACGGACG

ACCTGGCTCTGGGTTGTCTCCCTCACGGTGCACAAGCTTACCCCGCACAC

TTCCTCGACCGCCTTACGGTCGAGAAAAGCGAGGACTATATGCCCTCGCA

AACTAGCATGGCTAAATCGGACCCCAATAGCAATGACCTCCGGCAGGATC

GATTTTCGTTGCTGCACGGTCCACACGAGTTCTCACCCGTGCTTCCACCC

CCGCCGTTGACGGGTCCTTCGTCCCATTGTAATGGGTGTTCAGATACCCG

AGTGACCGTACGAGTCCTTGCGGATTTGCGGTCACCTATGTTGTTACTAG

ACACCCCCAATAGATAGCAGCCGACCTGTCTCACGACGGTCTAAACCCAG

ATGAGGGCGAGTTTCAGCCCTCAATCCGAACTACGATCGAGTTTAGGAGA

CGGTCGCTCTACCTCACGAACTATCTCATCAGAGGTCATGCTTCGACATG

CGACGCCTCGAACCTCGGCCGGTAAGCTGTTACGCTATTCTTAGAGGGTA

TTAGATGCGTTCAGCTCTTACCCCGTGGTGCGTAGCTGCCCGGCATCTGC

ACCTCCCGGCTGTTTAGGGCTCGAGACAATCTTCAATCACACTTAATCCA

ACGGTACGAGCTGACGGCGGCCATGCACCTCCTCTCTACAGCGTCGTGAT

GAGTGTTTAGTCTTCGCAGTTGATGCCTGCGAAATTCACGAGGGATATCC

GAGTCACTGCGACCTGCTCGGTCAAGAGCAGGCATCCCTTATCGCGAACT

AGTTCTTTCGTCGAGTGACCTTGTATGCCACGTCGCTCTATCGCCAACTG

CATTCATGCAAGCCGCTACTGAAGCGGCAAGGTACTACGCTACCTTAAGA

GAGATAGCAACCCATTGTCTCGACCATTGTAGCCCGCGTGTAGCCCAGCT

CTCGCGTGAGGAAATCTTTGGGCGCGCTCGATATCTTTTCGAGCGCGTAC

GTTTGGACTGTGTCGCGTTTACTCGCCGTTACTAACGACATCGCGGATTG

TATCGGTCATCACTTGAGCTGCCGGTATTACCGCGGCGGCTGGCACCGGT

CCAACTCTAACGGGCCTCCACGCAGCTTTCGCCACGCTTCGCCCTGCTCC

CCGAGGAACCCTTGCGCGTAAGGCCGTCGGGGTTCACACCCGACTCTCGC

CTCCGGTTGTAGTGCTCCCCCGCCAATTCCTTTAAGTTTCATCCTTGCGG

GTACACCAGTGGCACCCATTCGTAGTTCCTCTCGTACTATACGAACGTTC

TCGGAGATCCTCAGTTCTTTCCCTCCATGCGGGTCCCTGAGGCTTATCGC

GAGACTTCACATAAGACCCAGGTCCGTTATGGGTAGTCCGTACACCACAT

TCATCGGCTCTCGAGCCGAGCTATCCACCAGCTGGCACAGTAGCCAGTTG

CCCACCGTCTACCTAATCGGCCGCAGCCACATCCTACAGCGCCTGAGCGT

AGTTTGGCCACGTGTTACTGAGCTTTTCGCCACGAGTATGAACTCGTGCA

CCGAATTCCCTAACGTCGGTTGATCCCGACAGGCCTTGGCTTGCTCTGCC

CCCTTTCATCTAATTCGAATTACGGTTAGACTTAGGATCGACTAACCCTC

TTCCAGCATGACTCCCGTATGAACTATTAGCCTCAGTTTCCCGAGGTTAT

GGAACAACCCCGGGATCTGGTTCTTATGTTACGGGGTTTTCACCCTGTAT

GGCTCGCTTCTCGGCTTCCCTACGGCACAACACAGGCTCGTAGCCTGTGT

CAGATTCCGTCTCCGGGCTCTTGCTCTCACAACCCGTACCGATTATCGGC

GACACTCGTTGGGAATGTCACTGTCAGACTTCCGTTTGCTCTTGCGCTCT

TCGCACTTGCGTGCAGTGTAAAGGTTTCGCGCCTGCTGCGCCCCGTAGGG

CCCAGATTCGACCACCGTGTGGTGGCCTCATCCGGACCTCACTCGGGTGC
